# New *SNCA* mutation and structures of α-synuclein filaments from juvenile-onset synucleinopathy

**DOI:** 10.1101/2022.11.23.517690

**Authors:** Yang Yang, Holly J. Garringer, Yang Shi, Sofia Lövestam, Xianjun Zhang, Abhay Kotecha, Mehtap Bacioglu, Atsuo Koto, Masaki Takao, Maria Grazia Spillantini, Bernardino Ghetti, Ruben Vidal, Alexey G. Murzin, Sjors H.W. Scheres, Michel Goedert

**Author notes:** MOE Frontier Science Center for Brain Science and Brain-machine Integration, Zhejiang University, Hangzhou, China. Equal contributions (Yang Yang, Holly J. Garringer, Yang Shi).

## Abstract

A 21-nucleotide duplication in one allele of *SNCA* was identified in a previously described disease with abundant α-synuclein inclusions that we now call juvenile-onset synucleinopathy (JOS). Both wild-type α-synuclein and its insertion mutant containing seven additional residues (MAAAEKT) after residue 22 were present in sarkosyl-insoluble material that was extracted from frontal cortex of the individual with JOS and examined by electron cryo-microscopy. The structures of JOS filaments, comprising either a single protofilament, or a pair of protofilaments, revealed a new α-synuclein fold that differs from the folds of Lewy body diseases and multiple system atrophy (MSA). The JOS fold consists of a compact core, the sequence of which (residues 36-100 of wild-type α-synuclein) is unaffected by the mutation, and two disconnected density islands (A and B) of mixed sequences. There is a non-proteinaceous cofactor bound between the core and island A. The JOS fold resembles the common substructure of MSA type I and type II dimeric filaments, with its core segment approximating the C-terminal body of MSA protofilaments B and its islands mimicking the N-terminal arm of MSA protofilaments A. The partial similarity of JOS and MSA folds extends to the locations of their cofactor-binding sites. Our findings provide insight into a likely mechanism of JOS fibrillation in which mutant α-synuclein of 147 amino acids forms a nucleus with the JOS fold, around which wild-type and mutant proteins assemble during elongation.

## INTRODUCTION

α-Synuclein is the major component of the filamentous inclusions found in several neurodegenerative diseases, including Parkinson’s disease (PD), dementia with Lewy bodies (DLB) and multiple system atrophy (MSA) (9). We showed previously that assembled α−synuclein adopts distinct molecular conformers in the MSA and Lewy folds (25, 36). Most cases of PD and DLB are sporadic, but some are inherited in a dominant manner. Gene dosage and missense mutations have been described, with mean ages of onset of disease in the fourth and fifth decades (4,11,21,27,37).

Some of us previously described a case of synucleinopathy with an age of onset of disease of 13 years and death two years later that we named early-onset DLB (31). The proband developed rapidly progressing parkinsonism and cognitive impairment. Abundant α-synuclein-positive pathology was present throughout the cerebral cortex and several subcortical nuclei. Severe neuronal loss and gliosis in cerebral cortex and substantia nigra were present alongside vacuolar changes in the upper layers of the neocortex. Lewy pathology exceeded that of typical DLB.

We now report the cryo-EM structures of α-synuclein filaments from the frontal cortex of this case. They are different from the Lewy fold observed in PD and DLB, and share only a partial similarity with the structures of MSA filaments (25). Since distinct α-synuclein folds characterise different diseases (25, 36), this unique case appears to represent a distinct disease, which we now name juvenile-onset synucleinopathy (JOS).

We also report the likely cause of JOS: a previously overlooked 21- nucleotide duplication in the second exon of one allele of *SNCA*. This mutation, translating into insertion of the seven-residue sequence MAAAEKT after residue 22 of α-synuclein, probably occurred *de novo*. Both wild-type and mutant proteins are present in the JOS filaments. Unravelling their structures allowed us to propose a likely mechanism of fibrillation, in which the filaments are nucleated by the longer mutant α- synuclein and elongated through incorporation of both wild-type and mutant proteins.

## MATERIALS AND METHODS

### Clinical history and neuropathology

The case with JOS has been reported (31). A 13-year-old individual developed rapidly progressive parkinsonism, cognitive impairment, dysarthria and myoclonus, and died two years later. The number of α- synuclein deposits exceeded that of typical DLB.

### Sequencing of *SNCA* coding exons

Genomic DNA was extracted from frontal cortex. Coding exons of *SNCA* and flanking intronic sequences were amplified by polymerase chain reaction, screened by agarose gel electrophoresis and sequenced using the dideoxy method. Amplified exon 2 DNA from the JOS case was subcloned into pCR2.1 TA (Invitrogen). Recombinant plasmids were isolated from 48 clones and DNA from 5 of these clones was sequenced in both directions, as described (34).

### Extraction of α-synuclein filaments

Sarkosyl-insoluble material was extracted from frontal cortex of the individual with JOS, as described (32). In brief, tissues were homogenised in 20 vol (v/w) extraction buffer consisting of 10 mM Tris-HCl, pH 7.5, 0.8 M NaCl, 10% sucrose and 1 mM EGTA. Homogenates were brought to 2% sarkosyl and incubated for 30 min at 37° C. Following a 10 min centrifugation at 10,000g, the supernatants were spun at 100,000g for 20 min. Pellets were resuspended in 500 μl/g extraction buffer and centrifuged at 3,000g for 5 min. Supernatants were diluted threefold in 50 mM Tris-HCl, pH 7.5, containing 0.15 M NaCl, 10% sucrose and 0.2% sarkosyl, and spun at 166,000g for 30 min. Sarkosyl-insoluble pellets were resuspended in 100 μl/g of 20 mM Tris-HCl, pH 7.4.

### Immunolabelling, histology and silver staining

Immunogold negative-stain electron microscopy and immunoblotting were carried out as described (8). Filaments were extracted from the frontal cortex of the individual with JOS. PER4, a rabbit polyclonal serum that was raised against a peptide corresponding to residues 116-131 of human α-synuclein (29), was used at 1:50. Images were acquired at 11,000x with a Gatan Orius SC200B CCD detector on a Tecnai G2 Spirit at 120 kV. For immunoblotting, samples were resolved on 4-12% Bis-Tris gels (NuPage) and primary antibodies were diluted in PBS plus 0.1% Tween 20 and 5% non-fat dry milk. Before blocking, membranes were fixed with 1% paraformaldehyde for 30 min. Primary antibodies were: Syn303 [a mouse monoclonal antibody that recognises residues 1-5 of human α-synuclein (7)] (BioLegend) at 1:4,000, Syn1 [a mouse monoclonal antibody that recognises residues 91-99 from the NAC region of human α-synuclein (20)] (BD Biosciences) at 1:4,000 and PER4 at 1:4,000. Histology and immunohistochemistry were carried out as described (24, 26). Following deparaffinisation, the sections (8 μm) underwent heat-induced epitope retrieval in 10 mM citrate buffer, pH 6.0. The primary anti-α-synuclein antibodies were: Syn1 (1:250), 4B12 [a mouse monoclonal antibody that recognises residues 103-108 of human α-synuclein (12)] (BioLegend) at 1:250 and MJFR14 [a rabbit monoclonal antibody that has been reported to recognise assembled human α- synuclein (15)] (Abcam) at 1:1,000. Anti-Iba1 (Antibodies.com) and anti-GFAP antibodies (Dako), which recognise microglia and astrocytes, respectively, were used at 1:200. For signal detection, we used the protocols provided with the ImmPress HRP anti-rabbit/mouse IgG and ImmPACT SG kits (Vector Laboratories) or Alexa fluorophore-conjugated seconday antibodies (Invitrogen) at 1:250. Luminescent conjugated oligothiophene pFTAA (pentameric form of formyl thiophene acetic acid) was used at 3 μM, as described (14). To visualise inclusions, sections were also silver-impregnated according to Gallyas-Braak (2, 6). Some sections were counterstained with nuclear fast red (Vector Laboratories).

### Expression and purification of recombinant α-synuclein

Wild-type and mutant human α-synuclein was expressed and purified using modifications of a published protocol (16, 19). Constructs were made via cloning assembly using human α-synuclein in pRK172.

Expression was carried out in *E. coli* BL21(DE3) cells that were induced with 1 mM IPTG at an OD_600_ of 0.8 for 3-4 h and centrifuged at 20,000 g at 4° C for 20 min. They were resuspended in cold buffer (10 mM Tris-HCl, pH 7.4, 5 mM EDTA, 0.1 mM AEBSF, supplemented with cOmplete EDTA-free protease inhibitor cocktail) and lysed by sonication using a Sonics VCX-750 Vibra Cell Ultrasonic processor, followed by centrifugation at 20,000 g at 4° C for 45 min. The pellets were discarded and the pH of the supernatants was lowered to 3.5 with HCl, stirred for 30 min at room temperature and centrifuged at 50,000 g at 4° C for 1 h. The pH of the supernatants was then increased to 7.4 with NaOH and they were loaded onto an ion exchange HiTrap Q HP column and eluted with a 0-1 M NaCl gradient. Fractions containing α-synuclein were precipitated with ammonium sulphate (0.3 g/ml) for 30 min at 4° C and centrifuged at 20,000 g at 4° C for 30 min. The pellets were resuspended in PBS and loaded onto a HiLoad 16/60 Superdex (GE Healthcare) column equilibrated in PBS. The purity of α-synuclein was analysed by SDS-PAGE and protein concentrations were determined spectrophotometrically using an extinction coefficient of 5,600 M^-1^ cm^-1^. The protein-containing fractions were pooled and concentrated to 3-6 mg/ml. To separate wild-type and mutant α-synuclein, the samples were run on 16% Tricine gels (Invitrogen) and immunoblotted with Syn1.

### Electron cryo-microscopy

Extracted filaments were centrifuged at 3,000g for 3 min and applied to glow-discharged holey carbon gold grids (Quantifoil Au R1.2/1.3, 300 mesh), which were glow-discharged with an Edwards (S150B) sputter coater at 30 mA for 30 s. Aliquots of 3 μl were applied to the grids and blotted for approximately 3-5 s with filter paper (Whatman) at 100% humidity and 4° C, using a Vitrobot Mark IV (Thermo Fisher). Datasets were acquired on a Titan Krios G4 microscope (Thermo Fisher Scientific) operated at 300 kV. Images were acquired using a Falcon-4i detector and a Selectris-X energy filter (Thermo Fisher Scientific) with a slit width of 10 eV to remove inelastically scattered electrons. Images were recorded with a total dose of 40 electrons per Å^2^.

### Helical reconstruction

Movie frames were gain-corrected, aligned, dose-weighted and summed into a single micrograph using RELION’s own motion correction program (40). Contrast transfer function (CTF) parameters were estimated using CTFFIND-4.1 (38). Subsequent image-processing steps were performed using helical reconstruction methods in RELION (10,23,41). Filaments were picked manually. Reference-free 2D classification was performed to select suitable segments for generating initial models and for further processing. Initial models were generated *de novo* from 2D class averages using *relion helix inimodel2d* (23). Helical twist and rise were optimised during 3D autorefinement. Bayesian polishing and CTF refinement were used to further improve the resolution of reconstructions (41). Final maps were sharpened using standard post-processing procedures in RELION and their resolutions were calculated based on the Fourier shell correlation (FSC) between two independently refined half-maps at 0.143 (22). Helical symmetry was imposed on the post-processed maps using the *relion helix toolbox* program.

### Model building

Atomic models were built in Coot (3) using a shared substructure of MSA Type II filament (PDB:6XYQ) as template. Coordinate refinements were performed using *Servalcat* (35). Final models were obtained using refinement of only the asymmetric unit against the half-maps in *Servalcat*.

## RESULTS

### Genetic analysis

PCR amplification of control exon 2 DNA of *SNCA* produces a fragment of 377 nucleotides. Gel electrophoresis of amplified exon 2 DNA from the JOS patient revealed the presence of two fragments, the wild-type band and an additional band of 398 nucleotides (Figure 1a). DNA sequencing showed a 21-nucleotide duplication (TGGCTGCTGCTGAGAAAACCA) in one allele of *SNCA* (c.47_67dup). In α-synuclein, this translates into the insertion of seven residues (MAAAEKT) after residue 22, resulting in a protein of 147 amino acids. The PCR products of exon 2 amplification from JOS were subcloned into pCR2.12 and DNA sequencing confirmed the presence of both wild-type and mutant alleles. The mutation probably occurred *de novo*, since the proband’s parents did not exhibit symptoms of parkinsonism or cognitive impairment.

**Figure 1.**
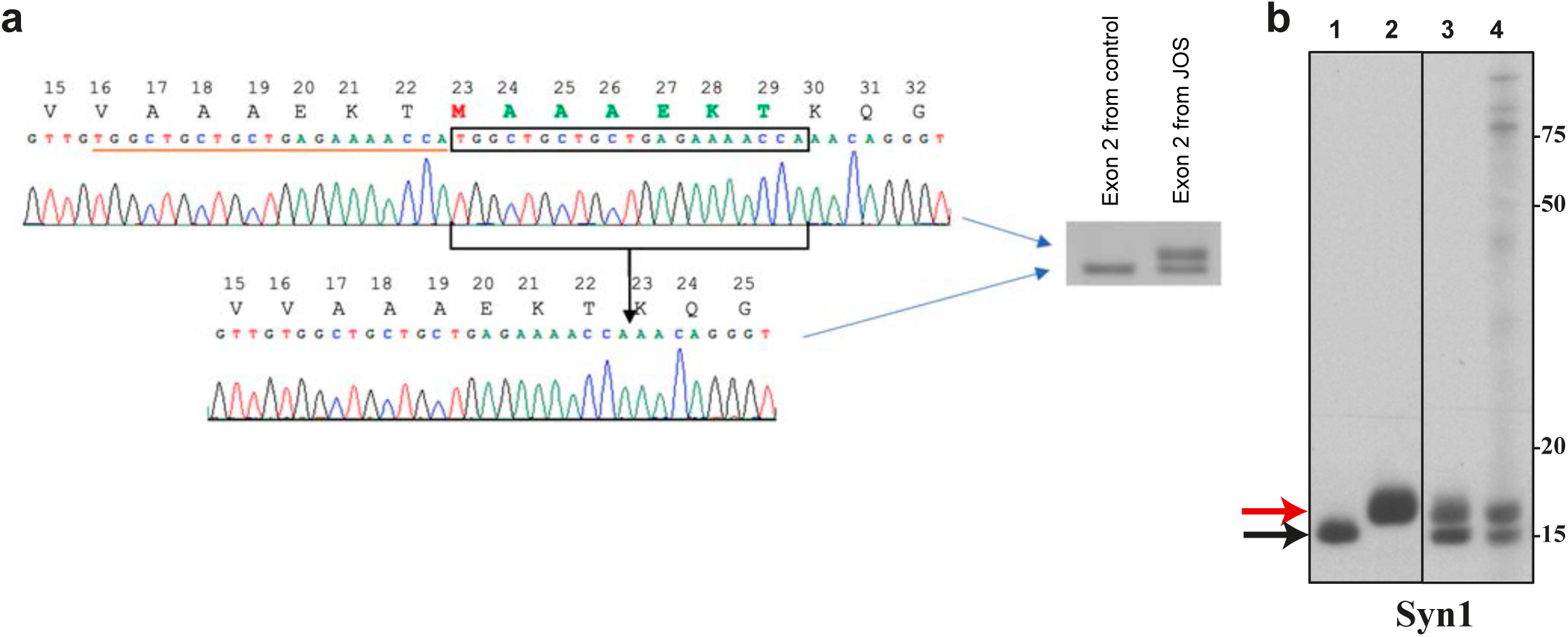
Seven amino acid insertion mutation in JOS α-synuclein. a, PCR amplification of *SNCA* exon 2 DNA from JOS gave the wild-type band of 377 nucleotides and a band of 398 nucleotides. Sequencing of the latter showed the presence of a 21 bp duplication, resulting in an in-frame insertion of seven amino acids (MAAAEKT) in the predicted protein. b, Immunoblotting of recombinant wild-type and mutant α-synuclein, and of JOS sarkosyl-insoluble material using Syn1. Lanes: 1, recombinant wild-type α-synuclein; 2, recombinant mutant α-synuclein; 3, mixture of recombinant wild-type and mutant α-synuclein; 4, JOS sarkosyl-insoluble material.

### Immunoblotting of sarkosyl-insoluble material

Recombinant wild-type (140 amino acids) and mutant (147 amino acids) α-synuclein ran as discrete bands of approximately 15 kDa and 16 kDa on 16% Tricine gels. When mixed together, they gave two separate bands on immunoblots with Syn1. Immunoblots of sarkosyl-insoluble material from the frontal cortex of the JOS individual gave two separate bands of similar intensities that aligned with the bands of recombinant wild-type and mutant α-synuclein (Figure 1b).

### Neuropathology

In cerebral cortex and several subcortical nuclei of JOS, we previously reported the presence of abundant intracytoplasmic α-synuclein inclusions in nerve cell bodies and processes (31). These findings were confirmed by immunostaining of frontal cortex with anti-α-synuclein antibodies Syn1, 4B12 and MJFR14 (Figure S1a). The α-synuclein inclusions of JOS were also strongly labelled by luminescent conjugated oligothiophene pFTAA (Figure S1b) and were weakly Gallyas-Braak silver-positive (Figure S1c). Triple-labelling immunofluorescence using anti α-synuclein antibody Syn1, as well as anti-Iba1 and anti-GFAP antibodies, showed prominent gliosis in areas with α-synuclein inclusions (Figure S1d). By immunoelectron microscopy of sarkosyl-insoluble material, abundant α-synuclein filaments with a diameter of 10 nm were present (Figure S2a). By immunoblotting with antibodies SYN303, SYN1 and PER4 (Figure S2b), high-molecular weight material, as well as full-length α-synuclein were the predominant species. Truncated α-synuclein was less abundant. None of the antibodies distinguished between wild-type and mutant α-synucleins.

### Structures of α-synuclein filaments from JOS

The vast majority of α-synuclein filaments from the frontal cortex of the individual with JOS (83%) comprise a single protofilament. The structure of these filaments was determined at 2.0 Å resolution and revealed a new α-synuclein fold that we call the JOS fold. In the structure of the remaining α-synuclein filaments (17%), determined at 2.3 Å resolution, there are two identical protofilaments with the JOS fold that pack against each other with C2 symmetry (Figure 2; Figure S3). Both singlet and doublet filaments have a left-handed helical twist, as established from the densities for backbone oxygen atoms in the cryo-EM maps (Figure S4).

**Figure 2.**
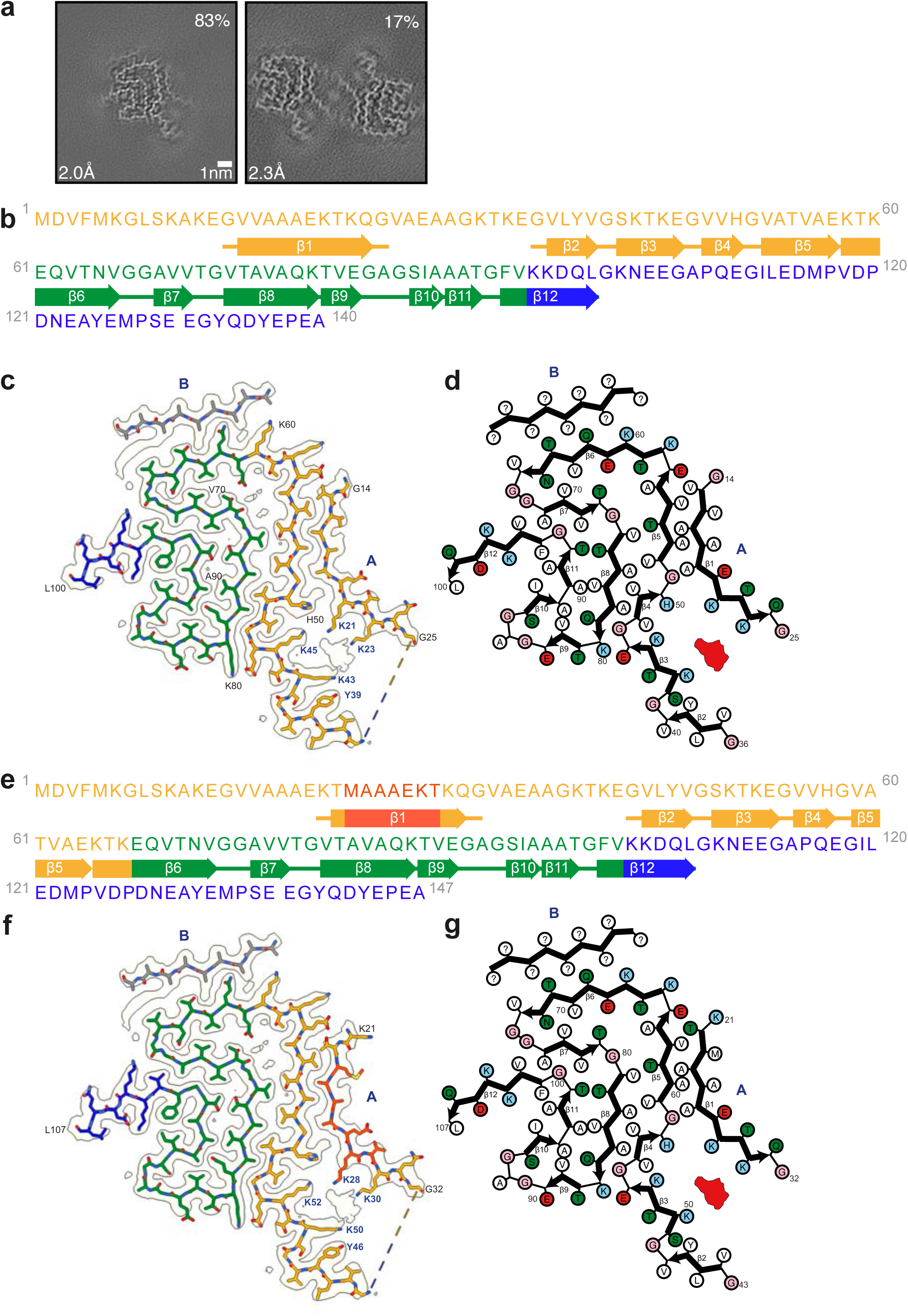
Cryo-EM structures of α-synuclein filaments from JOS. a, Cross-sections through the cryo-EM reconstructions, perpendicular to the helical axis and with a projected thickness of approximately one rung, are shown for α-synuclein singlet and doublet filaments from JOS. Scale bar, 1 nm. b,e, Amino acid sequences of wild-type (b) and mutant (e) α-synucleins. The N-terminal regions (1-60 or 1-67) are shown in yellow, the non-amyloid component regions (residues 61-95 or 68-102) in green and the C-terminal regions (residues 96-140 or 103-147) in blue. The insert (MAAAEKT) is indicated in red. Thick connecting lines with arrowheads indicate β-strands. c,f, Unsharpened cryo-EM density map and atomic model of the JOS fold with wild-type (c) or mutant (f) α-synuclein. Island B is indicated in grey. Potent cofactor densities are shown in red. d,g, Schematics of JOS fold with wild-type (d) or mutant (g) α-synuclein. Negatively charged residues are in red, positively charged residues in blue, polar residues in green, non-polar residues in white, sulfur-containing residues in yellow and glycines in pink. Thick connecting lines with arrowheads indicate β-strands. Unknown residues are indicated by question marks. Potent cofactor densities are shown in red.

The JOS fold consists of a compact core, spanning residues 36-100 (in the numbering of wild-type α-synuclein), and two disconnected density islands (A and B). There is also a large non-proteinaceous density between the core and island A that may correspond to a bound cofactor and is similar in size and environment to the unidentified densities of Lewy and MSA folds (25, 35). An additional density, next to the side chain of H50, may represent a post-translational modification. Other densities probably correspond to solvent molecules.

The core sequence is unaffected by the mutation and it has virtually the same conformation for residues K43-Q99 as the corresponding segment of MSA protofilament IIB2 (PDB:6XYQ) (25). The conformation of this segment, which is stabilised in part by a characteristic salt bridge between residues E46 and K80, is also similar to the conformations of wild-type α-synuclein filaments following seeded aggregation using MSA brain seeds (PDB:7NCK) (16) and H50Q (PDB:6PEO) (1) or A53T (PDB:6LRQ) (28) α-synuclein filaments assembled *in vitro*, with minor differences in the inversion of side chain orientations of residues K58 and T59 (Figure 3).

**Figure 3.**
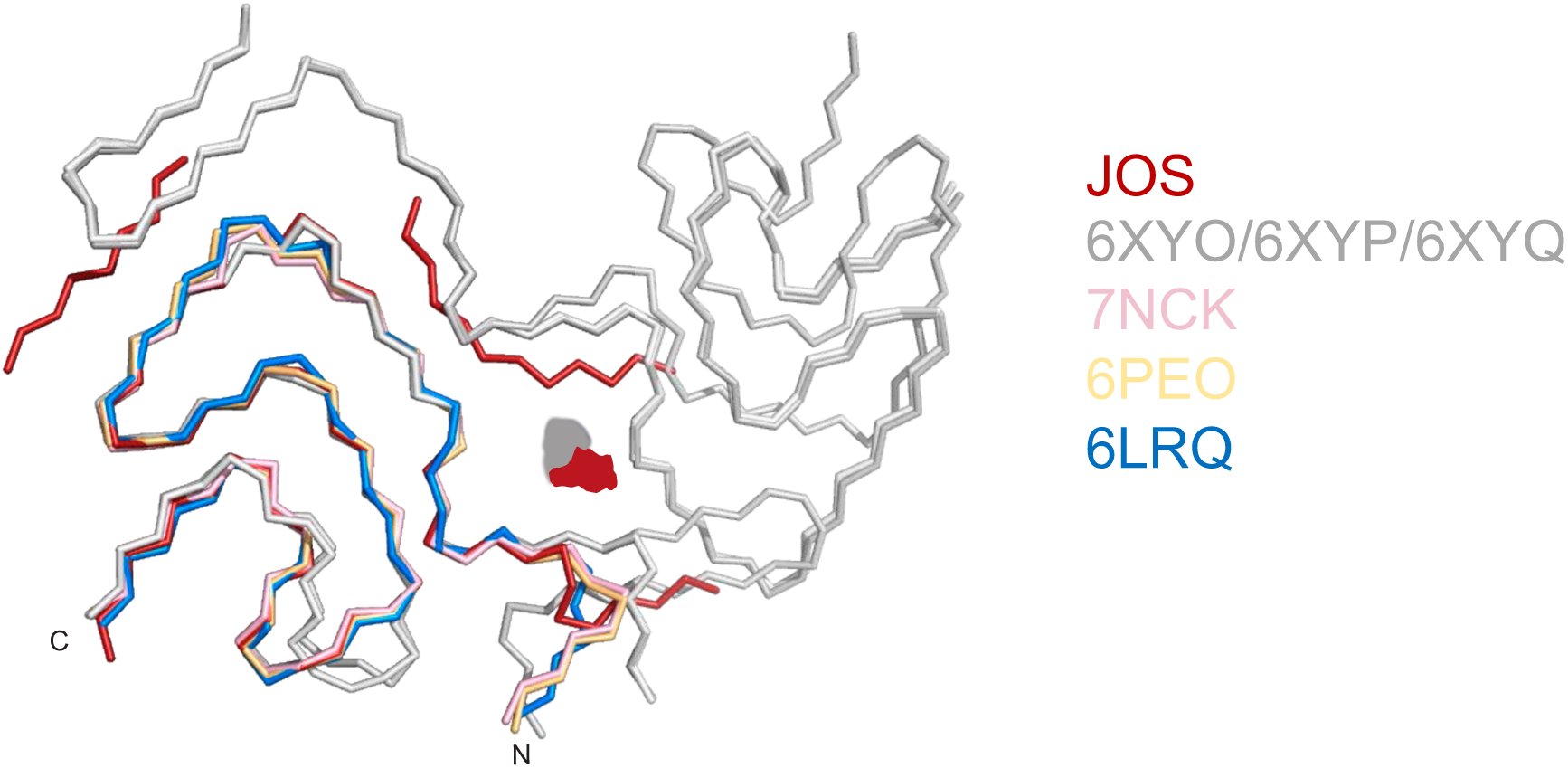
Comparison of α-synuclein folds. Ribbon plot of the JOS fold in red; MSA folds in grey (6XYO/6XYP/6XYQ); seeded assembly using MSA brain seeds in pink (7NCK); assembled recombinant H50Q α-synuclein in yellow (6PEO); assembled recombinant A53T α-synuclein in blue (6LRQ). Potential co-factor densities are shown in red for JOS and in grey for MSA.

In the core, most side chain densities are fully resolved in the 2.0 Å resolution map of JOS singlet filaments. In the islands, however, the side chain densities are not well resolved, precluding unambiguous sequence assignment. Since their main chain densities are fully resolved, we attribute the lack of distinct side chain densities to the islands consisting of a mixture of different sequences. These sequences probably come mainly from the amino- and/or carboxy-terminal regions of the same α- synuclein chain, but incorporation of exogenous peptide fragments cannot be excluded.

In the structure of island A, there are at least three clues that point to which sequences are incorporated. Island A makes a curved β-strand, the amino-terminal half of which is tightly packed against residues A53 and V55 of the core, whereas its carboxy-terminal half bends away opposite residue H50 and faces the non-proteinaceous density, separating it from residues K43 and K45. Its interface with the core resembles part of the protofilament interface of MSA filaments between the N-terminal arm of protofilaments A and the C-terminal body of protofilaments B. The MSA protofilament interface harbours a non-proteinaceous density that is surrounded by four lysine residues in approximately the same location, suggesting that the residues in island A that face a similar density in the JOS fold are also lysines. A residue with a small side chain, which is packed between A53 and V55, is almost certainly alanine. Side chain densities for Cβ atoms strongly suggest that island A comprises ten consecutive non-glycine residues.

Only two α-synuclein sequences satisfy all three clues: _14_GVVAAAEKTKQG_25_ in wild-type protein (Figure 2c-d) and _21_KTMAAAEKTKQG_32_ in the 147-residue insertion mutant (Figure 2f-g).

There are insufficient clues to suggest which sequences the shorter island B is made of. However, we do note that our assignment of island A with the insertion mutant leaves an N-terminal segment that is long enough to reach and fill up the island B density. A model of the JOS fold made only of mutant protein, in which island B is assigned the sequence _6_KGLSKAKE_13_ and is connected to island A (_21_KTMAAAEKTKQG_32_), is shown in Figure S5. In this model, residues 13-20, which connect islands B and A, mimic the conformation of the N-terminal arm of MSA protofilaments A. The absence of density for these residues in the cryo-EM map indicates that they are not ordered in the JOS filaments. An alternative explanation for the density of island B is that it can be reached from the last ordered residue of the core (L100) and that it consists of sequences from the C-terminal region of α-synuclein.

The cryo-EM reconstructions of JOS doublet filaments show that island A is essential for dimerisation, with the islands from both protofilaments facing each other across the filament axis. The 2.3 Å resolution map of the doublet filament further supports our sequence assignments for island A. Interestingly, there are no direct contacts between the ordered residues of both protofilaments. Like in dimeric filaments of TMEM106B (26), an additional disconnected density of unknown identity resides on the symmetry axis and mediates their interactions (Figure S3). This non-proteinaceous density, which is co-ordinated by neutral polar residues Q24 (Q31 in mutant α-synuclein) from both protofilaments, is probably of a different chemical nature than the large non-proteinaceous densities within each protofilament. The density of island B, which is farthest from the doublet filament axis, is less resolved than in the singlet filament map, but is similar in overall shape.

## DISCUSSION

We report a duplication of 21 nucleotides in one allele of *SNCA* in a case of JOS, which inserts seven amino acids (MAAAEKT) after residue 22 of α-synuclein. This is the first duplication mutation in *SNCA*. It was not present in the 141,456 individuals of the gnomAD project (13). The clinical picture was more severe than in DLB and the age of onset (13 years) was atypical. Previously, missense mutations in the coding region of *SNCA* were reported in inherited cases of PD and DLB (21, 37). Gene dosage mutations (duplications and triplications) of *SNCA* have also been described (4,11,27). Typically, individuals with these mutations have a family history of synucleinopathy. The individual with JOS had no family history, suggesting that the mutation occurred *de novo*.

The presence of extensive nerve cell loss and abundant filamentous α- synuclein inclusions (31) was confirmed and the latter were weakly Gallyas-Braak silver-positive. α-Synuclein inclusions of MSA are strongly Gallyas-Braak silver-positive, whereas those of PD and DLB are negative (33). Luminescent conjugated oligothiophene pFTAA strongly labelled α- synuclein inclusions from the frontal cortex of the JOS individual, similar to what has been described in Parkinson’s disease and multiple system atrophy (14). As reported previously (31), by negative stain immuno-EM, these filaments had the same staining characteristics as those of PD, DLB and MSA (5,29,30). However, by cryo-EM, a different fold was present.

Immunoblotting of sarkosyl-insoluble material showed that both wild-type and mutant α-synucleins make up the JOS filaments. It is conceivable that mutant α-synuclein forms a nucleus, around which wild-type and mutant proteins assemble during elongation, giving rise to mixed filaments. This is reminiscent of mutation R406W in tau, where almost equal amounts of wild-type and mutant tau were present in the sarkosyl-insoluble fractions (18).

The JOS fold is the third known fold of assembled α-synuclein from human brain, the other two being the MSA and Lewy folds (Figure 4) (25, 36). The JOS fold differs from the Lewy fold, but it resembles the common substructure of MSA type I and type II filaments, with its core segment adopting nearly the same structure as the C-terminal body of MSA protofilaments B and their islands mimicking the N-terminal arms of MSA protofilaments A. This similarity of JOS and MSA folds extends to the locations of their cofactor-binding sites. In the brain of the JOS case, almost all a-synuclein inclusions were present in neurons, indicating that a fold similar to MSA folds can form in nerve cells. In the core part, the JOS fold is similar to the folds of recombinant α-synuclein filaments (1,16,28). However, there is a small, but probably essential, difference between the JOS fold and the recombinant α-synuclein filament folds in one turn at K58. In the recombinant α-synuclein filament folds, this lysine side chain points inside the core, whereas it points outside in the JOS and MSA folds. In the MSA dimeric filaments, K58 of protofilament B makes a salt bridge with E35 of protofilament A. There is a second salt bridge between K60 of protofilament B and E28 of protofilament A.

**Figure 4.**
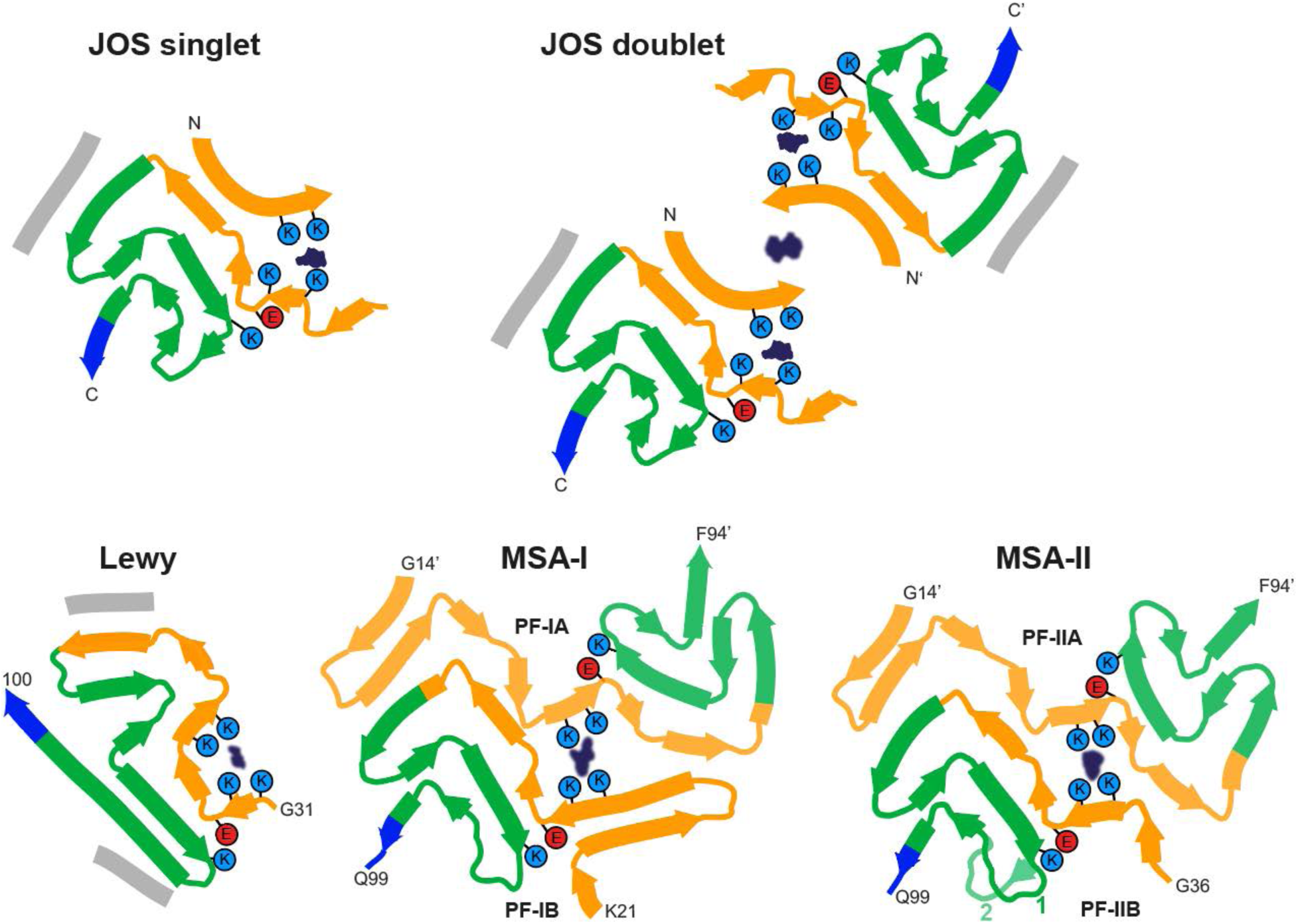
JOS, Lewy and MSA folds of α-synuclein from human brains. Schematics of secondary structure elements in JOS, Lewy and MSA folds, depicted as single rungs, and coloured as in Figure 2. Potential co-factor densities are indicated in dark blue. The positions of their surrounding residues, as well as the supporting salt bridges between E46 and K80 in JOS and MSA folds, and between E35 and K80 in the Lewy fold, are highlighted with coloured circles. Residue numbers with apostrophes indicate those from the other protofilament.

In our model of JOS filaments that are only made of mutant α- synuclein, there are equivalent salt bridges between K65 and E20, and between K67 and E13, which stabilise the interaction between the filament core and the N-terminal region. This implies that mutant α- synuclein would be prone to form filament nuclei, possibly affected by the presence of a cofactor, that then could grow by incorporation of both wild-type and mutant proteins.

## ACKNOWLEDGEMENTS

This work was supported by the Electron Microscopy Facility of the MRC Laboratory of Molecular Biology. We thank Jake Grimmett, Toby Darling and Ivan Clayson for help with high-performance computing. We acknowledge Diamond Light Source for access and support of the cryo-EM facilities at the UK’s Electron Bio-Imaging Centre (under proposal bi23268), funded by the Wellcome Trust, the MRC and the Biotechnology and Biological Sciences Research Council (BBSRC). M.G. is an associate member of the UK Dementia Research Institute.

## AUTHOR CONTRIBUTIONS

H.J.G., M.B., A.K., M.T., M.G.S., B.G. and R.V. identified the patient and performed genetic analysis and neuropathology. Y.Y. prepared filaments, performed immunoblots and immunoelectron microscopy; X.Z. and A.K. performed cryo-EM data acquisition; Y.Y., Y.S., A.G.M. and S.H.W.S. performed cryo-EM structure determination; S.H.W.S. and M.G. supervised the project and all authors contributed to the writing of the manuscript.

## DECLARATIONS

### Ethics approval and consent

Studies carried out at Indiana University were approved through the Institution’s ethical review process. Informed consent was obtained from the patient’s next of kin. This study was approved by the Cambridgeshire 2 Research Ethics Committee (09/H0308/163) and by Mihara Memorial Hospital (085–01).

### Data and materials availability

Cryo-EM maps have been deposited in the Electron Microscopy Data Bank (EMDB) with the accession numbers EMD-16188 and EMD-16189. Corresponding refined atomic models have been deposited in the Protein Data Bank (PDB) under accession numbers 8BQV and 8BQW. Please address requests for materials to the corresponding authors.

## SUPPLEMENTARY FIGURE LEGENDS

**Figure S1.**
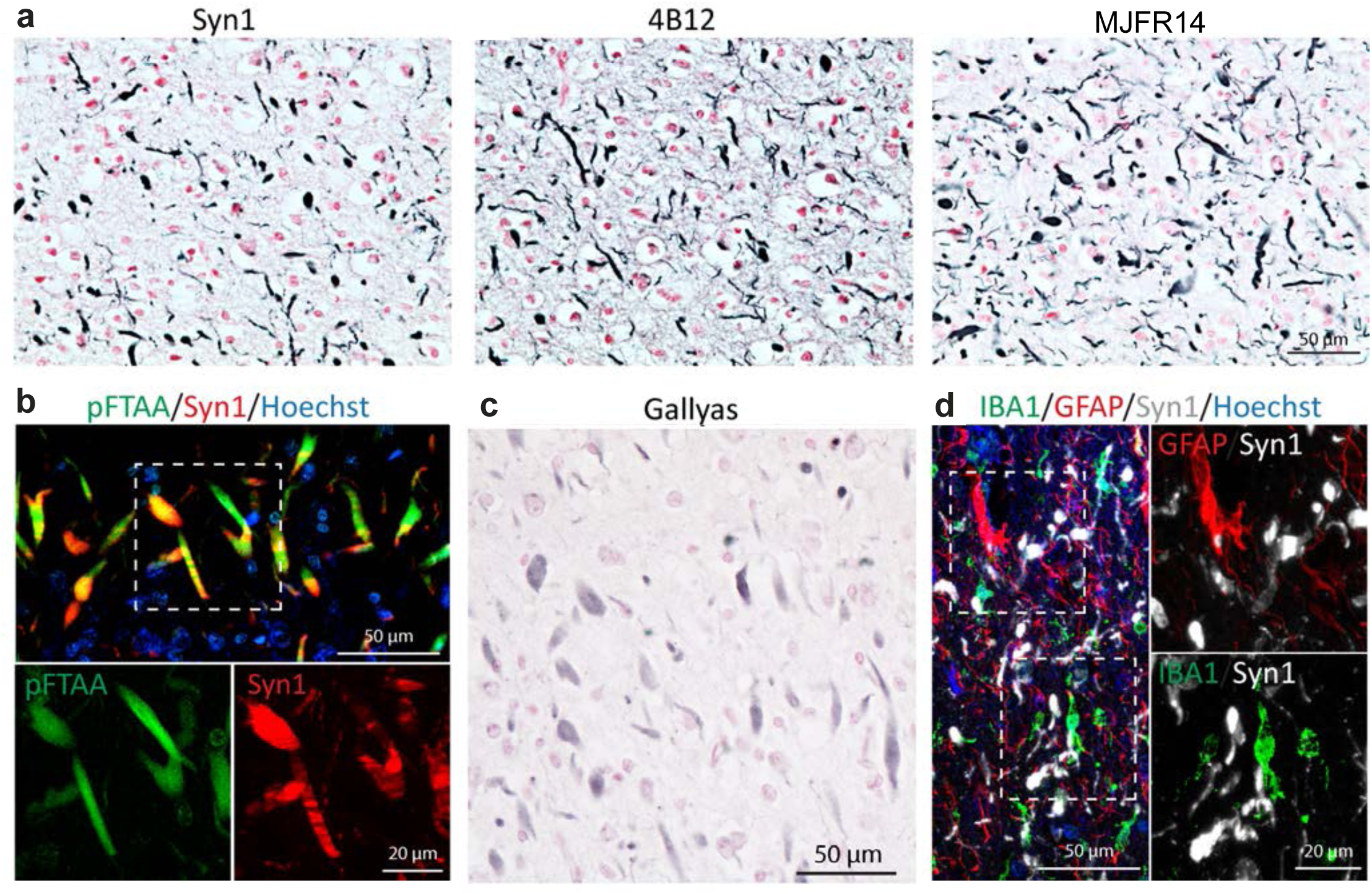
Staining of α-synuclein inclusions from frontal cortex of JOS. a, Immunostaining for α-synuclein using antibodies Syn1, 4B12 and MJFR14, showing numerous stained inclusions (blue). Nuclei are stained red. b, Double-labelling immunofluorescence using anti-α-synuclein antibody Syn1 (red) and luminescent conjugated oligothiophene pFTAA (green). Double labelling (yellow). Nuclei are stained blue (Hoechst dye). c, Gallyas-Braak silver staining (black). Nuclei are stained red. d, Triple-labelling immunofluorescence using anti-α-synuclein antibody Syn1 (white), anti-microglial marker Iba1 (green) and anti-astrocyte marker GFAP (red). Nuclei are stained blue (Hoechst dye).

**Figure S2.**
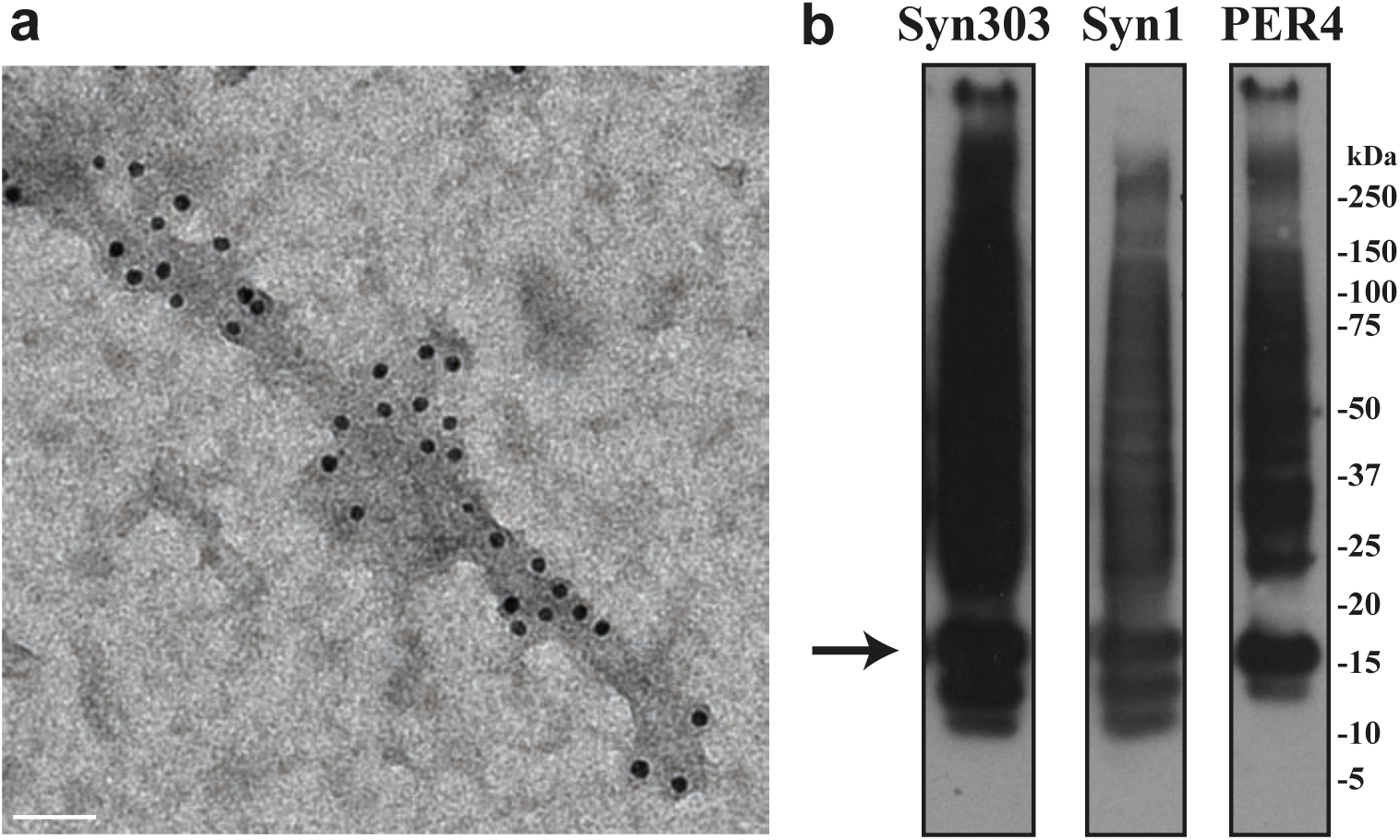
Negative-stain immunoelectron microscopy and immunoblotting of α-synuclein filaments from frontal cortex of JOS. a, Negative-stain immunoelectron microscopy of JOS filaments, with polyclonal antibody PER4. b, Immunoblotting of JOS filaments. Sarkosyl-insoluble material was blotted with monoclonal antibodies Syn303 and Syn1, and polyclonal antibody PER4. The arrow points to monomeric α-synuclein.

**Figure S3.**
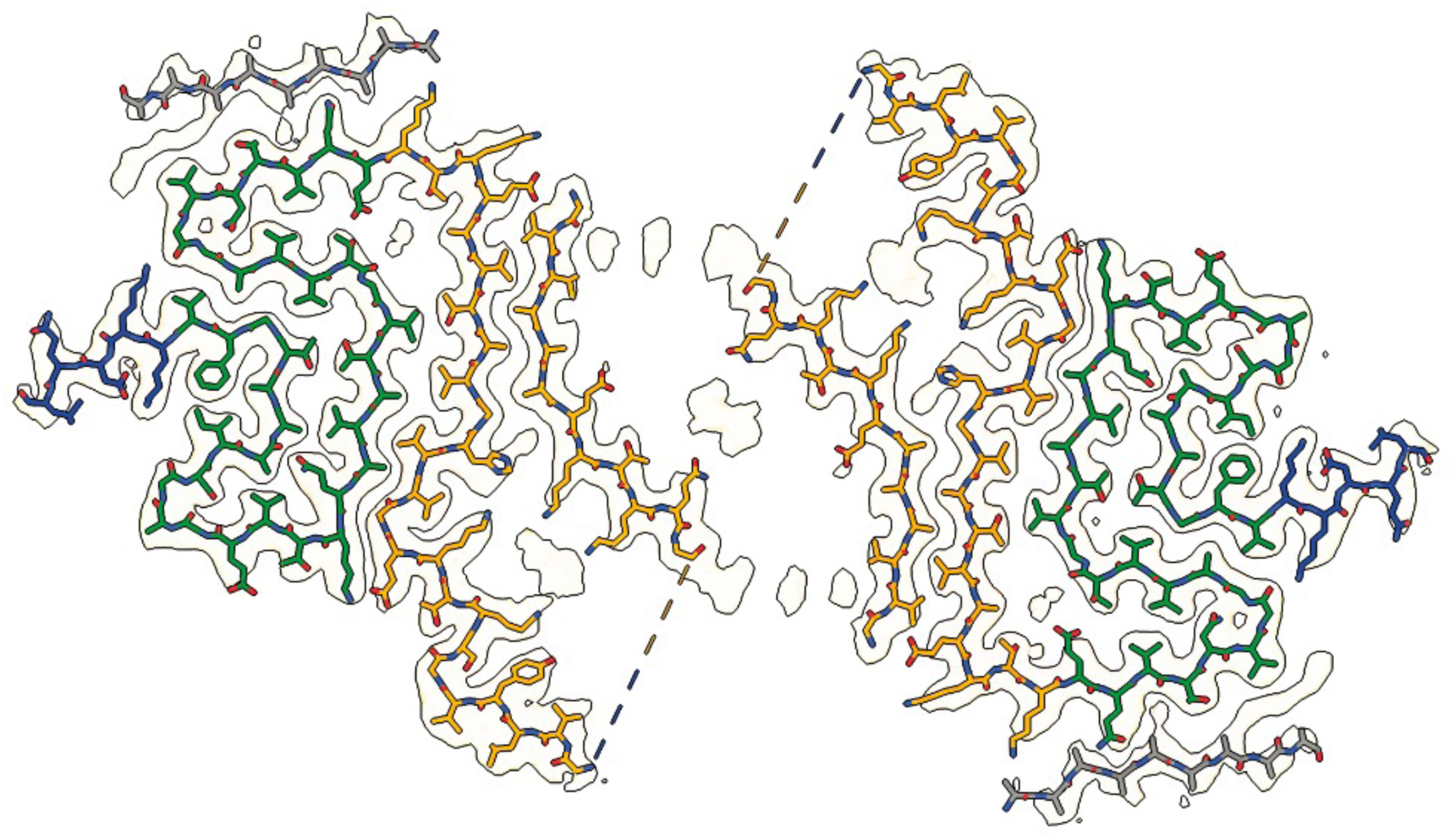
Cryo-EM structure of JOS doublet filaments. Cryo-EM density map and atomic model of JOS fold (wild-type) in doublet. The model is coloured as in Figure 2.

**Figure S4.**
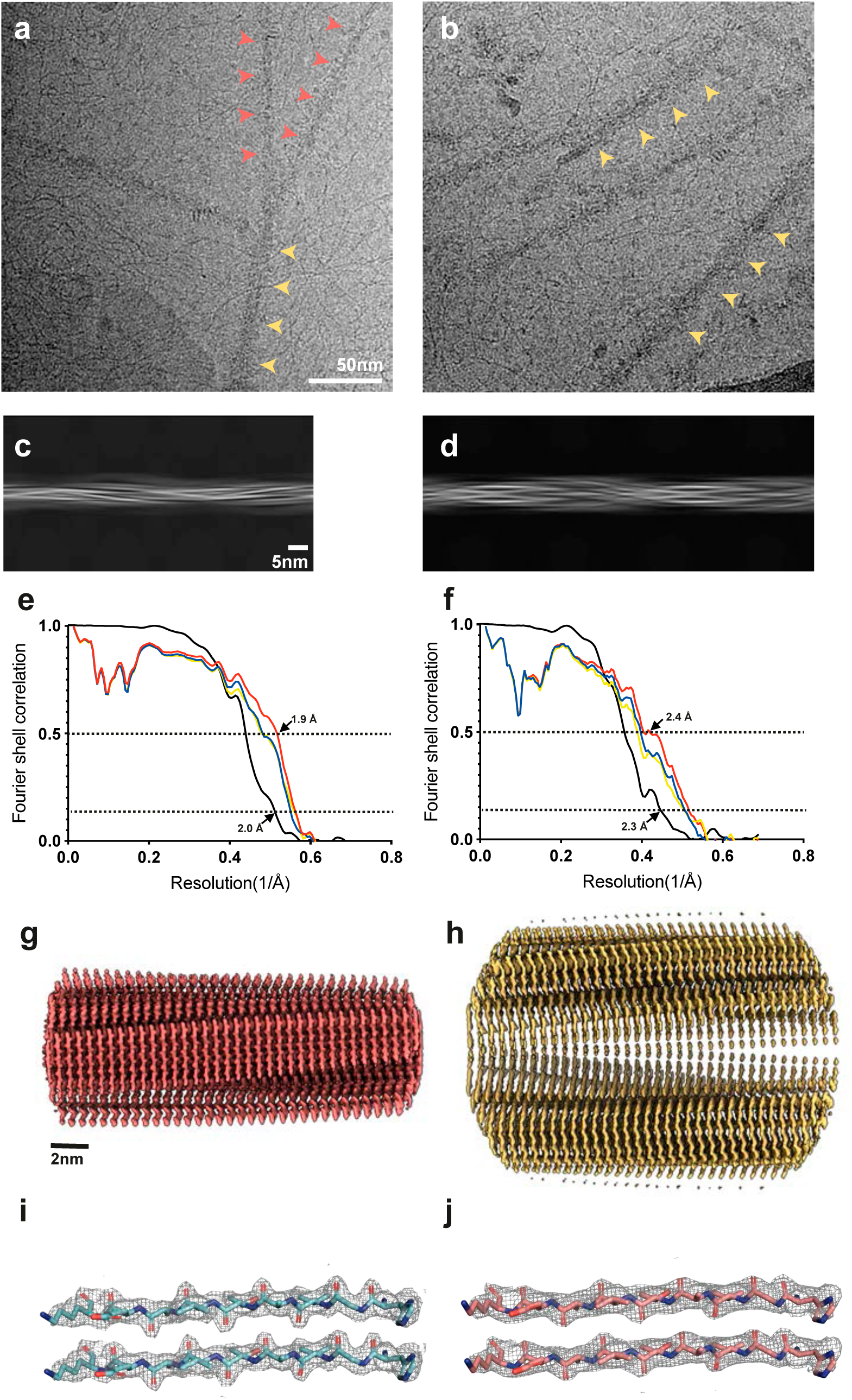
Cryo-EM images and resolution estimates. a,b, α-Synuclein filaments from JOS. Orange arrowheads (a) point to singlet, yellow arrowheads (a and b) to doublet JOS filaments. Scale bar, 50 nm. c,d, Projection features of singlet (c) and doublet (d) JOS filaments. Scale bar, 5 nm. e,f, Fourier shell correlation (FSC) curves for cryo-EM maps and structures of α-synuclein filaments in JOS singlet (e) and doublet (f) filaments. FSC curves for two independently refined cryo-EM half maps are shown in black; for the final refined atomic model against the final cryo-EM map in red; for the atomic model refined in the first half map against that half map in blue; for the refined atomic model in the first half map against the other half map in yellow. g,h, Side views of JOS singlet (g) and doublet (h) filaments. i,j, Zoomed-in views of the main chains of singlet (i) and doublet (j) filaments, showing the densities of main-chain oxygen atoms.

**Figure S5.**
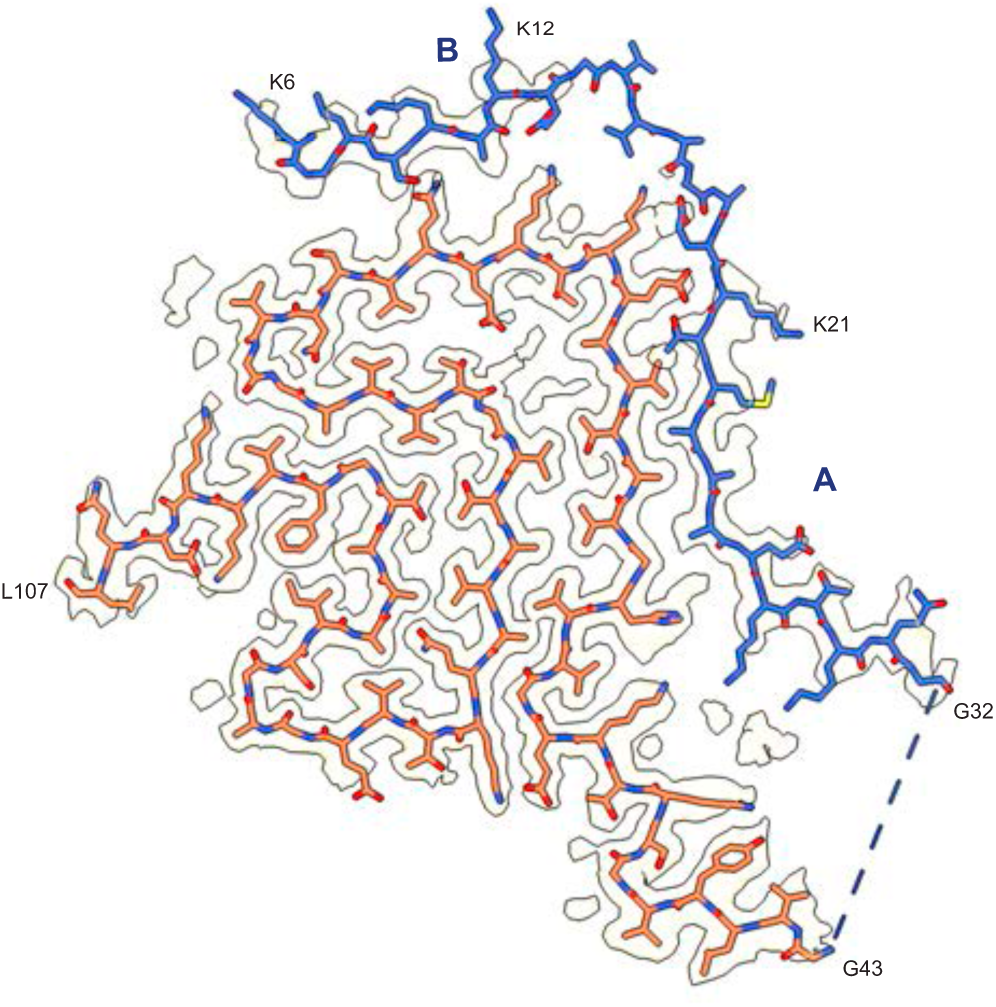
Model of the JOS fold with mutant α-synuclein. Island B is assigned the sequence _6_KGLSKAK_12_ and connected to island A (_21_KTMAAAEKTKQG_32_), both in blue. Core regions are in orange and map densities in grey.

**Table S1.** Cryo-EM data acquisition and structure determination.

|  | JOS fold (singlet)<br>PDB 8BQV<br>EMD-16188 | JOS fold (doublet)<br>PDB 8BQW<br>EMD-16189 |
| --- | --- | --- |
| **Data acquisition** |  |  |
| Electron gun | CFEG | CFEG |
| Detector | Falcon 4i | Falcon 4i |
| Energy filter slit (eV) | 10 | 10 |
| Magnification | 165,000 | 165,000 |
| Voltage (kV) | 300 | 300 |
| Electron dose (e<sup>-</sup>/Å<sup>2</sup>) | 40 | 40 |
| Defocus range (μm) | 0.6 to 1.4 | 0.6 to 1.4 |
| Pixel size (Å) | 0.727 | 0.727 |
| **Map refinement** |  |  |
| Symmetry imposed | C1 | C2 |
| Initial particle images (no.) | 354269 | 354269 |
| Final particle images (no.) | 214769 | 45038 |
| Map resolution (Å) | 2 | 2.3 |
| FSC threshold | 0.143 | 0.143 |
| Helical twist (°) | -1.33 | -1.11 |
| Helical rise (Å) | 4.77 | 4.71 |
| **Model Refinement** |  |  |
| Model resolution (Å) | 2 | 2.3 |
| FSC threshold | 0.5 | 0.5 |
| Map sharpening *B* factor (Å<sup>2</sup>) | -33 | -43 |
| Model composition |  |  |
| Non-hydrogen atoms | 1719 | 3477 |
| Protein residues | 252 | 516 |
| Ligands | 0 | 0 |
| *B* factors (Å<sup>2</sup>) |  |  |
| Protein | 35.5 | 43.9 |
| R.m.s. deviations |  |  |
| Bond lengths (Å) | 0.0025 | 0.0068 |
| Bond angles (°) | 0.64 | 1.27 |
| Validation |  |  |
| MolProbity score | 1.82 | 1.64 |
| Clashscore | 7.20 | 3.39 |
| Poor rotamers (%) | 0 | 0 |
| Ramachandran plot |  |  |
| Favored (%) | 93.59 | 91.25 |
| Allowed (%) | 6.41 | 8.75 |
| Disallowed (%) | 0 | 0 |

